# Dredd-mediated cleavage of Kenny uncouples the IKK complex from selective autophagy to enable innate immunity

**DOI:** 10.64898/2026.04.24.720600

**Authors:** Aravind K. Mohan, Anna M. Dahlström, Anna L. Aalto, Kerttu Kotala, Veera Luukkonen, Fanny Serenius, Elinda Helin, Tor Erik Rusten, Annika Meinander

**Affiliations:** Faculty of Science and Engineering, Biochemistry and Cell Biology, Åbo Akademi University, Turku, Finland; InFLAMES Research Flagship Center, Åbo Akademi University, Turku, Finland; Department for Clinical Medicine, Medical Faculty, University of Oslo, Department of Molecular Cell Biology, Institute for Cancer Research, Oslo University Hospital, Oslo, Norway

**Author notes:** **Correspondence:** Annika Meinander, Faculty of Science and Engineering, Biochemistry and Cell Biology, Åbo Akademi University, Tykistökatu 6, 20520 Turku. Equal contribution (Aravind K. Mohan, Anna M. Dahlström).

**Keywords:** Autophagosome, Caspase, *Drosophila*, Dredd, IKK, Imd, Kenny, NF-κB, Ref(2)P

## Abstract

Selective autophagy restrains innate immune signalling to maintain tissue homeostasis, yet how this repression is rapidly relieved during infection remains unclear. Here, we show that under basal conditions the inhibitor of κB kinase γ (IKKγ) Kenny is sequestered at autophagosomes through Atg8 and the selective autophagy receptor Ref(2)P, thereby silencing Imd pathway activity. Bacterial infection disrupts this interaction, releasing the IKK complex to enable immune signalling. Mechanistically, we identify the initiator caspase Dredd as a direct interactor of the IKKγ Kenny and show that Dredd binds and cleaves Kenny in a ubiquitination-dependent manner during infection. This cleavage removes an N-terminal LC3-interacting region, uncoupling the IKK complex from autophagosomal degradation. Dredd-mediated processing of Kenny stabilises the IKK complex and is required for activation of the NF-κB transcription factor Relish, robust antibacterial responses, and host survival following infection. Together, these findings uncover a mechanism by which caspase-mediated cleavage intersects with selective autophagy to dynamically control NF-κB signalling during bacterial infection.

**Short summary:** Bacterial infection activates NF-κB signalling by triggering caspase-dependent cleavage of the IKK subunit Kenny, releasing the IKK complex from autophagosomal repression to enable effective innate immune responses.

## Introduction

Autophagy is an evolutionary conserved process that maintains cellular homeostasis. In macroautophagy (hereafter referred to as autophagy), cytoplasmic cargo is engulfed by a double membrane isolation membrane that upon fusion with the lysosome degrades and recycles cytoplasmic components. Autophagy was initially thought to be a non-discriminatory process that sequesters bulk cytoplasm. However, it has later been recognised that autophagy can selectively degrade cargo, such as organelles and proteins and is a key regulator of immunity and host defense^1^. The intestinal epithelium is an interface for host-microbe interactions and, therefore, an important immune organ. In *Drosophila*, the intestine is the first layer of defence against pathogens, and it functions both as a physical and a molecular barrier of local immune response reactions^2–5^. The fly intestine provides a convenient model to study local immune responses, as the intestinal epithelium can be exposed to both selected pathogens and compounds by feeding. By contrast, the fat body mediates systemic immune responses and is critical during acute infections, such as when pathogens breach epithelial barriers and gain access to the haemocoel^7,8,9^. Transcriptional activation of nuclear factor κ-light-chain-enhancer of activated B cells (NF-κB) transcription factors control the molecular immune responses, both in the gut and the fat body. The NF-κB transcription factor Relish is activated via the immune deficiency (Imd) pathway which is particularly important for local immune responses in the fly gut^3,7^. While the Imd pathway has been thoroughly studied, little is known about the role of how the stability of the signalling proteins is regulated by selective autophagy.

The *Drosophila* Imd pathway is primarily activated upon infection by Gram-negative bacteria^8,9^. Following receptor activation, a protein complex composed of Imd, Fas- associated death domain (Fadd), and the caspase-8 homologue, Death related ced-3/Nedd2-like protein (Dredd), is formed^12–14^. The E3 ligase *Drosophila* initiator of apoptosis 2 (Diap2) catalyses Lysine 63 (K63) ubiquitination of Imd and Dredd, which is required for activation of Dredd and the Imd pathway^16–18^. The inhibitor of κB kinase (IKK) complex is composed of a kinase subunit, IKKβ, termed Immune response deficient 5 (Ird5), and a regulatory subunit IKKγ, called Kenny, and it functions downstream of Dredd^19–21^. Kenny, like the mammalian IKKγ NF-κB essential modulator (NEMO), possesses a ubiquitin binding in ABIN and NEMO (UBAN) domain, through which it can bind Methionine 1 (M1) linked ubiquitin (Ub) chains^22^. The IKK complex has two important functions in activating Relish. Ird5 phosphorylates Relish, which is necessary for a robust transcriptional activation^24^ and the IKK complex is also required, independent of its kinase activity, for Dredd-mediated cleavage of Relish^19,23,24^. This activation of Relish culminates in production and secretion of a set of antimicrobial peptides (AMPs) for microbial control^7,10^.

Selective autophagy is mediated by cargo receptors that link ubiquitinated substrates to autophagosomes. Refractory to Sigma P (Ref(2)P), also called sequestome-1 (SQSTM1), binds ubiquitinated proteins and promotes their autophagic degradation through interaction with both ubiquitin and Autophagy-related 8 (Atg8) family proteins^26,45,46^. To prevent constitutive activation of the Imd pathway, Kenny has been shown to be selectively degraded by autophagy by interacting with autophagosomes through its microtubule-associated protein 1 light chain 3 (LC3) interacting region (LIR) motif^26^. Here, we have studied how the autophagosomal association and release of Kenny is regulated during basal conditions and infection and how this modulates immune signalling. We found that Ref(2)P interacts with Kenny during basal conditions, whereas bacterial challenge triggers binding of the caspase-8 homologue Dredd to Kenny. This interaction results in Dredd-dependent cleavage of the N-terminal LIR motif of Kenny. Removal of the LIR motif is required for Kenny to be released from Ref(2)P and the autophagosome and therefore being able to mount an intestinal immune response to Gram-negative bacteria. Hence, we describe a caspase-mediated mechanism of diversion of the IKK complex from autophagosomal degradation to being an active inducer of Relish activation.

## Results

### Bacterial infection releases Kenny and Ref(2)P from autophagosomes

The *Drosophila* IKK complex is a potent inducer of NF-κB activation via the Imd pathway. It has been shown that the IKK complex is targeted for autophagic degradation to prevent constitutive immune activation during basal conditions, and the interaction with the autophagosomes has been shown to be mediated by Atg8a and the LIR motif of *Drosophila* the IKKγ Kenny ^20,26^. To study autophagosome sequestration of Kenny in intestinal epithelial cells, we expressed N-terminally green fluorescent protein (GFP)-tagged Kenny in flies using the upstream activating sequence (UAS) Galactose-responsive transcription factor 4 (Gal4) system. Immunofluorescence analysis of the *Drosophila* midgut revealed that GFP-tagged Kenny can be found in the same cellular compartments as the autophagosome protein Atg8a and the selective autophagy receptor Ref(2)P (Supplementary Figure 1A). Feeding flies with chloroquine blocks the autophagic flux by hindering the fusion of lysosomes with autophagosomes. This impaired lysosomal degradation resulted in increased levels of both Kenny, Ref(2)P and Atg8a in the midgut (Figure 1A, Supplementary Figure 1B), which indicates that Kenny and Ref(2)P undergo autophagic degradation in the intestine during basal conditions.

**Figure 1.**
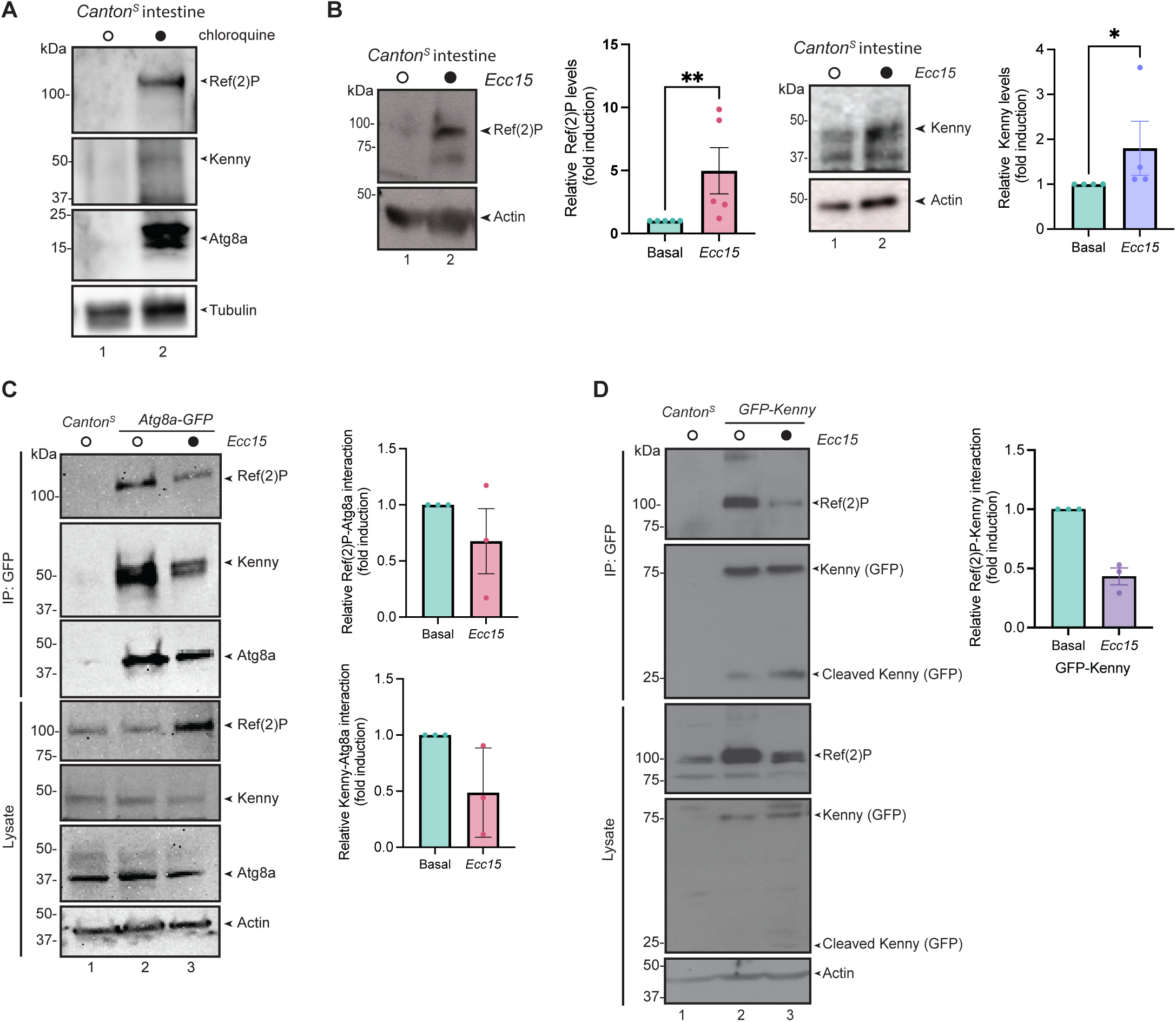
Ref(2)P and Kenny interact, and their interaction is reduced upon bacterial infection. **(A)** Adult Canton^S^ flies were fed with 5 % sucrose and 100 µM chloroquine for 16 h. Endogenous Ref(2)P, Kenny and Atg8a levels in lysates from adult intestines were analysed with Western blotting with anti-Ref(2)P, anti-Kenny, anti-Atg8a and anti-Tubulin antibodies, n=3. **(B)** Adult *Canton^S^* flies were fed with 5% sucrose and *Ecc15* for 16 h, Ref(2)P and Kenny levels were analysed in lysates from dissected intestines by Western blotting with anti-Ref(2)P, anti-Kenny, and anti-Actin antibodies. The relative protein levels of endogenous Ref(2)P and Kenny levels were quantified, n>4. **(C)** Adult *Canton^S^* and *Atg8a-GFP* flies were fed with 5% sucrose and *Ecc15* for 16 h before whole flies were lysed and GFP-immunoprecipitations were performed and the samples were analysed by Western blotting with anti-Ref(2)P, anti-Kenny, anti-Atg8a, and anti-Actin antibodies. The ratio of immunoprecipitated Ref(2)P and Kenny to total Ref(2)P and Kenny was quantified, n=3. **(D)** Adult wildtype *Canton^S^* and *GFP-Kenny* expressing flies were fed with 5% sucrose and *Ecc15* for 16 h before lysis. GFP-immunoprecipitations were performed and the samples were analysed by Western blotting with anti-Ref(2)P, anti-GFP, and anti-Actin antibodies. The ratio of immunoprecipitated Ref(2)P to total Ref(2)P was quantified, n=3.

To study autophagosomal association of Kenny and Ref(2)P during infection, we infected flies by feeding them with the Gram-negative bacteria *Erwinia carotovora* strain *Ecc15*. We observed increased Ref(2)P and Kenny levels upon infection in fly intestines, suggestive of defective autophagosomal turnover (Figure 1B, Supplementary Figure 2). To study the interaction of Kenny and Ref(2)P to the autophagosomal Atg8a, we immunoprecipitated GFP-tagged Atg8a from whole fly lysates. The results showed that endogenous Kenny and Ref(2)P co-precipitate with Atg8a during basal conditions, but that their interaction is reduced upon bacterial infection (Figure 1C). To analyse if Kenny and Ref(2)P interact, we co-immunoprecipitated GFP-tagged Kenny with Ref(2)P from fly lysates (Figure 1D) and lysates from S2 cells (Supplementary Figure 3). We found that Kenny and Ref(2)P interact with each other, but that their interaction was reduced upon infection by feeding flies with *Ecc15.* This suggests that Ref(2)P and Kenny are targeted to autophagosomes for degradation during basal conditions, and that their autophagosomal association and degradation is reduced upon infection when Kenny is required for Imd activation.

### Dredd interacts with and cleaves Kenny

When immunoprecipitating Kenny from fly lysates, we surprisingly observed an accumulation of a smaller fragment of Kenny after bacterial challenge, suggesting that infection induce cleavage of Kenny (Figure 1D). Dredd is an initiator caspase activated upon bacterial infection, and its catalytic activity is required for processing of signalling proteins to become active Imd pathway inducers^14,16^, and hence represent a potential protease for Kenny. To test if Kenny is a substrate for Dredd, we expressed Dredd and Kenny in S2 cells and found that co-expression with Dredd induces cleavage of Kenny (Figure 2A). By immunoprecipitating Kenny from S2 cell lysates, we found that Dredd and Kenny interact (Figure 2B). Dredd consists of a prodomain and a caspase domain (Figure 2C). While the caspase domain contains the p20 and p10 subunits harbouring the catalytic site, the prodomain contains two death effector domains (DED1 and DED2) that are important for recruitment to the complexes, in which Dredd is catalytically activated^27^. We found that Kenny interacts with the prodomain of Dredd (Figure 2D), and more specifically the DED1 (Figure 2E). As the cleaved fragment of C-terminally HA-tagged Kenny is approximately 45 kDa, we generated point mutations of all aspartic acid residues located in the N-terminal region of Kenny, i.e., D21E, D27E, D66E, D88E (Figure 2F). When co-expressing Dredd with these mutants in S2 cells, we found that wildtype and all Kenny mutants except D21E could be cleaved (Figure 2G) upon co-expression with Dredd, indicating that D21 is a Dredd cleavage site.

**Figure 2.**
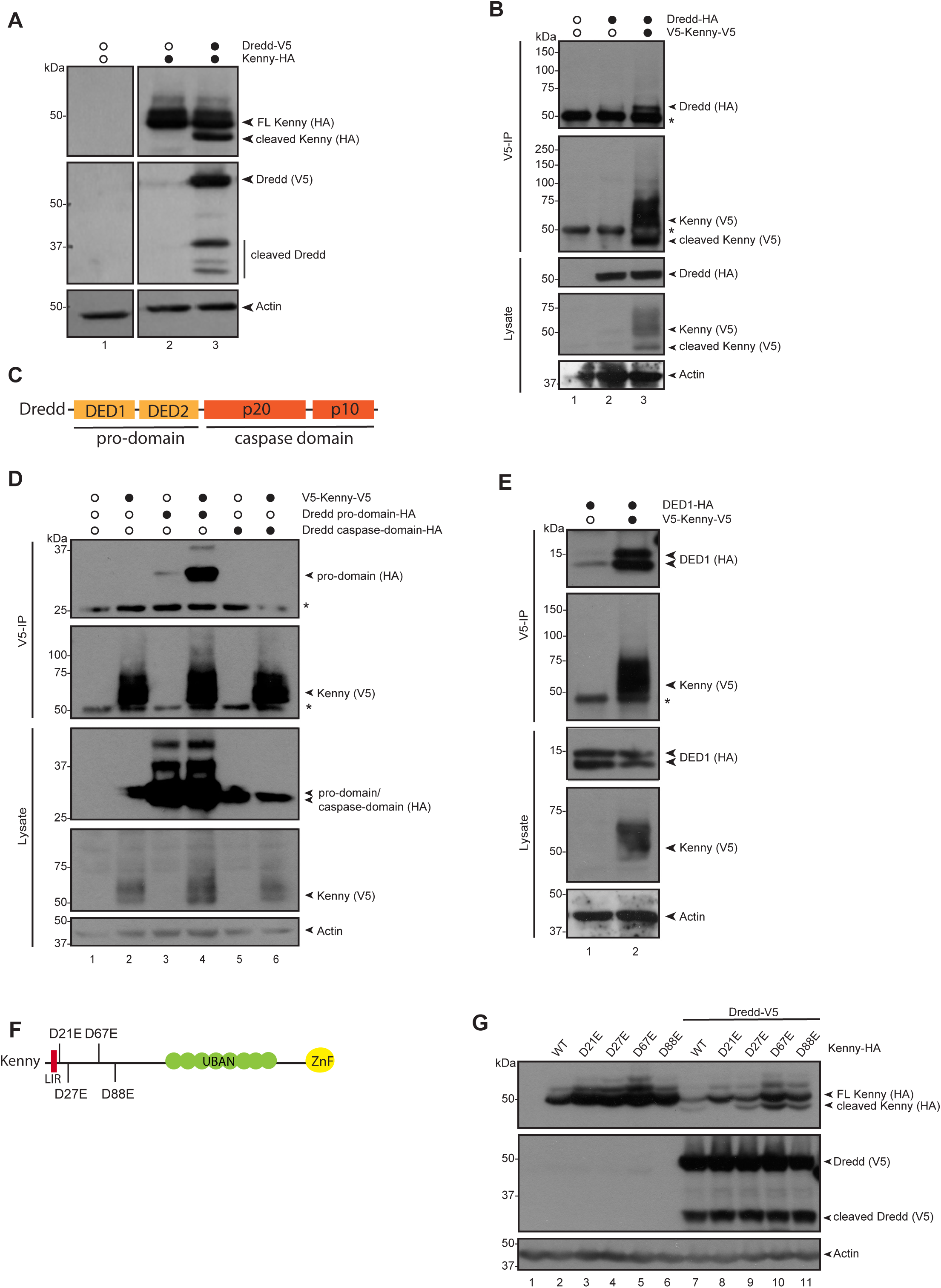
Dredd interacts with and cleaves Kenny. **(A)** *Drosophila* S2 cells were transfected with empty vector, V5-tagged Dredd, and HA-tagged Kenny. The cell lysates were analysed by Western blotting using anti-HA, anti-V5, and anti-Actin antibodies, n>3. **(B)** *Drosophila* S2 cells were transfected with empty vector, HA-tagged Dredd and V5-tagged Kenny. V5-immunoprecipitations were performed and the samples were analysed by Western blotting with anti-HA, anti-V5, and anti-Actin antibodies, n=3. **(C)** Schematic representation of Dredd. **(D)** *Drosophila* S2 cells were transfected with empty vector, V5-tagged Kenny, and HA-tagged pro-domain or caspase domain of Dredd. V5-immunoprecipitations were performed and the samples were analysed by Western blotting with anti-HA, anti-V5, and anti-Actin antibodies, n=3. (**E**) *Drosophila* S2 cells were transfected with V5-tagged Kenny, and HA-tagged DED1 of Dredd. V5-immunoprecipitations were performed and the samples were analysed by Western blotting with anti-HA, anti-V5, and anti-Actin antibodies, n=3. **(F)** Schematic representation of the HA-tagged Kenny construct showing the positions of D21E, D27E, D67E and D88E point mutations. **(G)** *Drosophila* S2 cells were transfected with empty vector, V5-tagged Dredd, and HA-tagged Kenny WT and HA-tagged Kenny mutants D21E/D27E/D67E/D88E. The cell lysates were analysed by Western blotting using anti-HA, anti-V5, and anti-Actin antibodies, n>3.

### Dredd-mediated Kenny cleavage requires Dredd ubiquitination

To analyse the interaction between DED1 of Dredd and Kenny we used molecular modelling. We based the modelling on the crystal structure of the complex of Kaposi’s sarcoma herpes virus (KSHV)-FLIP, which contains two DEDs similar to Dredd and mammalian Caspase-8 and NEMO, which is the mammalian IKKγ (PDB: 3CL3). We used the AlphaFold (Q8IRY7) model of Dredd (Figure 3A, blue) and aligned this structure onto the KSHV-FLIP dimer (Figure 3A, grey) bound to NEMO (Figure 3A, pink). The model suggests that the DED1 of Dredd can interact with IKKs similarly as the DED1 of KSVH-FLIP.

**Figure 3.**
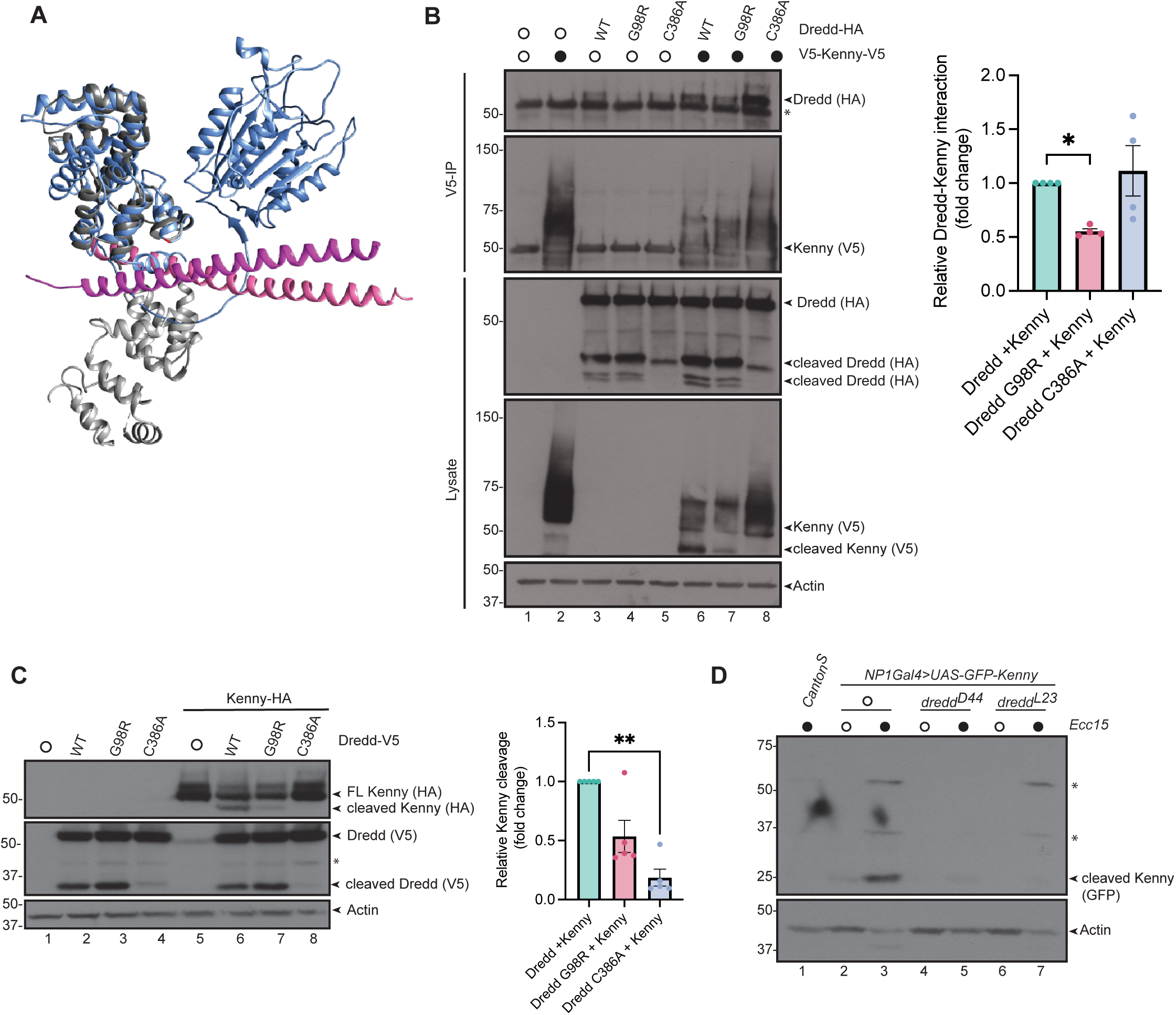
Kenny cleavage is dependent on Dredd’s DED1 domain. **(A)** Structural modelling of the Dredd-Kenny interaction. Dredd (blue, AlphaFold: Q8IRY7) is modelled on the complex of two KSHV-FLIP (grey) proteins associated with a NEMO (pink) dimer (PDB: 3CL3). The molecular graphics and analyses were performed with the UCSF Chimera package^41^. (**B**) *Drosophila* S2 cells were transfected with empty vector, HA-tagged wildtype, G98R, or C386A point mutant of Dredd and V5-tagged Kenny. V5-immunoprecipitations were performed and the samples were analysed by Western blotting with anti-HA, anti-V5, and anti-Actin antibodies. The ratio of immunoprecipitated Dredd to total Dredd was quantified, n=4. **(C)** *Drosophila* S2 cells were transfected with empty vector, V5-tagged wildtype, G98R, or C386A point mutant of Dredd and HA-tagged Kenny. Kenny cleavage was analysed from transfected S2 cells by Western blotting with anti-HA, anti-V5, and anti-Actin antibodies. The ratio of cleaved to total Kenny was quantified, n=5. **(D)** Adult male guts from *Canton^S^* flies or flies expressing GFP-Kenny (*NP1Gal4>UAS-GFP-Kenny*) in a wildtype, *dredd^D44^*, or *dredd^L23^* mutant background, fed with 5% sucrose and *Ecc15*, were dissected. Kenny cleavage was analysed from dissected guts lysed in lysis buffer and analysed by Western blotting with anti-GFP and anti-Actin antibodies, n=3.

Dredd has been shown to interact with Fadd and Diap2 through its DED1 domain, and this domain is also ubiquitinated by Diap2 during bacterial infection^16,28^. We have previously characterised a mutation in DED1 (G98R) of Dredd that affects its K63-ubiquitination by Diap2^16^. This point mutation was originally described as G120R, but the transcribed Dredd protein has later been reported to be 22 amino acids shorter than the original sequence. The ubiquitination-deficient G98R mutant Dredd showed significantly reduced interaction with Kenny compared to wildtype Dredd, whereas the catalytically inactive C386A mutant Dredd retained its interaction (Figure 3B). This demonstrates that Dredd can associate with Kenny independently of its catalytic activity, but Dredd ubiquitination strengthens this interaction. As expected, the reduced interaction of the G98R mutant Dredd to Kenny also reduced its ability to cleave Kenny (Figure 3C). These results suggest that DED1 interacts with Kenny, and that the cleavage of Kenny requires both ubiquitination and catalytic activity of Dredd.

As we had found that Kenny is cleaved upon bacterial infection, and that Dredd is able to cleave Kenny, we continued by analysing Dredd-mediated cleavage of Kenny during bacterial infection *in vivo*. For this purpose, we expressed GFP-tagged Kenny in the intestinal epithelium of flies using the NP1-Gal4 driver and fed the adult flies with the Gram-negative bacteria *Ecc15*. We were not able to detect Kenny in intestines from untreated flies, as Kenny is degraded during basal conditions. When expressing GFP-Kenny in wildtype flies, in *dredd^D44^*-mutant flies with reduced ubiquitination, or in *dredd^L23^*-mutant flies with reduced catalytic activity, we detected a robust infection-induced cleavage of Kenny in the presence of wildtype Dredd, but not in the Dredd mutants (Figure 3D). This suggests that loss of the catalytic activity of Dredd, either by catalytic mutation or by abrogation of ubiquitination, results in lack of processing of its target, Kenny. Thus, oral infection with *Ecc15* triggers the Kenny cleavage, protecting it from degradation.

### Dredd-mediated cleavage is required for stabilisation of Kenny

The LIR-domain of Kenny has been shown to be involved in targeting the IKK complex to selective autophagic degradation^25^. While this degradation is suggested to prevent constitutive activation of the Imd pathway during basal conditions^26^, the D21 site, where Kenny is cleaved by Dredd is localised just after the N-terminal region of Kenny that harbours the LIR-domain. To determine whether Dredd-mediated cleavage affects autophagosome association, we generated transgenic flies expressing a C-terminally HA-tagged wildtype Kenny and a cleavage-resistant Kenny mutant (D21E) regulated by the upstream activating element UAS. Immunofluorescence analysis of *Drosophila* midgut revealed that, under basal conditions both HA-tagged wildtype and D21E Kenny can be found in the same compartments as Atg8a and Ref(2)P. The D21E mutant Kenny is however accumulated in compartments colocalising with Ref(2)P and Atg8a compared to wildtype Kenny (Figure 4A, Supplementary Figure 4). To study the interaction between Kenny and the autophagy receptor Ref(2)P, we performed immunoprecipitations from whole adult fly lysates. We found that Ref(2)P co-immunoprecipitated with both wildtype and D21E mutant Kenny during basal conditions (Figure 4B, lane 3 and 5). While this interaction was modestly reduced during infection with *Ecc15* in flies expressing wildtype Kenny (Figure 4B), no reduction in the Kenny-Ref(2)P interaction was observed after bacterial infection in D21E Kenny mutants (Figure 4B). These results indicate that Kenny cleavage affects its interaction with Ref(2)P. Consequently, a strong induction of wildtype Kenny was detected in lysates after *Ecc15* infection, whereas no infection-induced upregulation of the uncleaved D21E mutant Kenny could be seen (Figure 4B, lane 4 and 6). Our results thus indicate that cleavage of Kenny upon infection promotes Kenny stabilisation.

**Figure 4.**
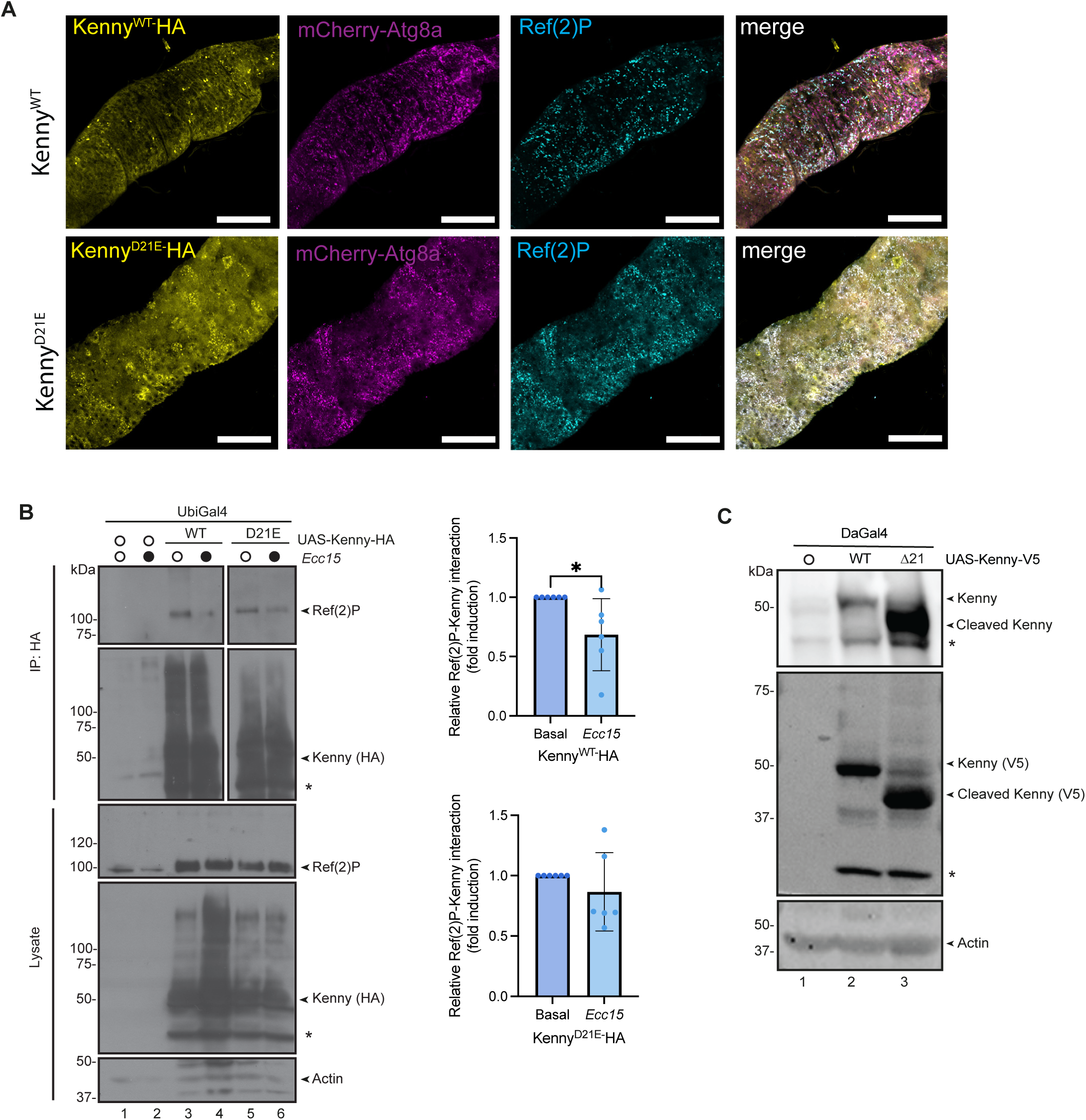
Dredd-mediated cleavage is required for stabilisation of Kenny. **(A)** Dissected intestines from adult flies expressing mCherry-Atg8a (magenta) crossed with flies expressing HA-tagged wildtype or D21E mutant Kenny were stained for HA (yellow) and Ref(2)P (cyan). Co-localisation of Atg8a, Kenny and Ref(2)P is indicated as white puncta in the merged image. Midguts were imaged by confocal microscopy using at least 3 intestines per repeat, n=3. Scale bar 100 µm. **(B)** Adult control *UbiGal4* flies and flies expressing HA-tagged wildtype or D21E mutant Kenny by control of UbiGal4 were fed with 5% sucrose and *Ecc15* for 16 h before lysis. HA-immunoprecipitations were performed and the samples were analysed by Western blotting with anti-Ref(2)P, anti-HA and anti-Actin antibodies, n=5. The relative interaction between Ref(2)P and Kenny were quantified. **(C)** Adult control *DaGal4* flies and flies expressing V5-tagged wildtype or Δ21 mutant Kenny by control of *DaGal4* were lysed and Kenny expression levels were analysed with Western Blotting with anti-Kenny (upper panel), anti-V5 (middle panel), and anti-Actin antibodies, n>3.

To further investigate whether Kenny cleavage affects its expression, we generated a constitutively cleaved Δ21 mutant Kenny lacking the LIR domain. We were able to detect both wildtype and constitutively cleaved Δ21 mutant Kenny during basal conditions in these flies. However, as expected, constitutively cleaved Kenny, which is not recognised by the autophagosome, is expressed more strongly compared to wildtype (Figure 4C). These results indicate that Kenny cleavage is crucial for stabilisation of Kenny by protecting it from autophagy-mediated degradation.

### Dredd-mediated cleavage of Kenny is required for NF-κB activation and immunity

To analyse whether Kenny cleavage is required for Relish-mediated activation of the Imd pathway in *Drosophila*, we examined the infection-induced expression of the Relish target gene *diptericin*. We expressed wildtype and mutant HA-tagged Kenny with the DaGal4 driver, which directs ubiquitous expression of UAS-regulated Kenny in a heterozygous Kenny loss-of-function (LOF) *key^4^-*mutant background. Upon *Ecc15* infection, we found that *diptericin* was induced in control flies and flies expressing wildtype Kenny. However, the ability of the cleavage-resistant D21E mutant Kenny to induce *diptericin* expression upon bacterial infection was impaired, similarly as LOF mutation of Kenny (Figure 5A). Concordantly, we found that expression of constitutively cleaved Kenny resulted in increased expression of the Relish target genes *diptericin* and *drosocin* under basal conditions (Figure 5B). These results indicate that Kenny cleavage is important for Relish activation and efficient immune responses during bacterial infection in flies. Likewise, we observed a significant increase in *diptericin* gene expression in Atg8a^RNAi^ flies both during basal conditions and *Ecc15* infection compared to control flies (Figure 5C). This suggests that Kenny cleavage is required for Relish activation and that autophagy normally serves to dampen the Imd pathway.

**Figure 5.**
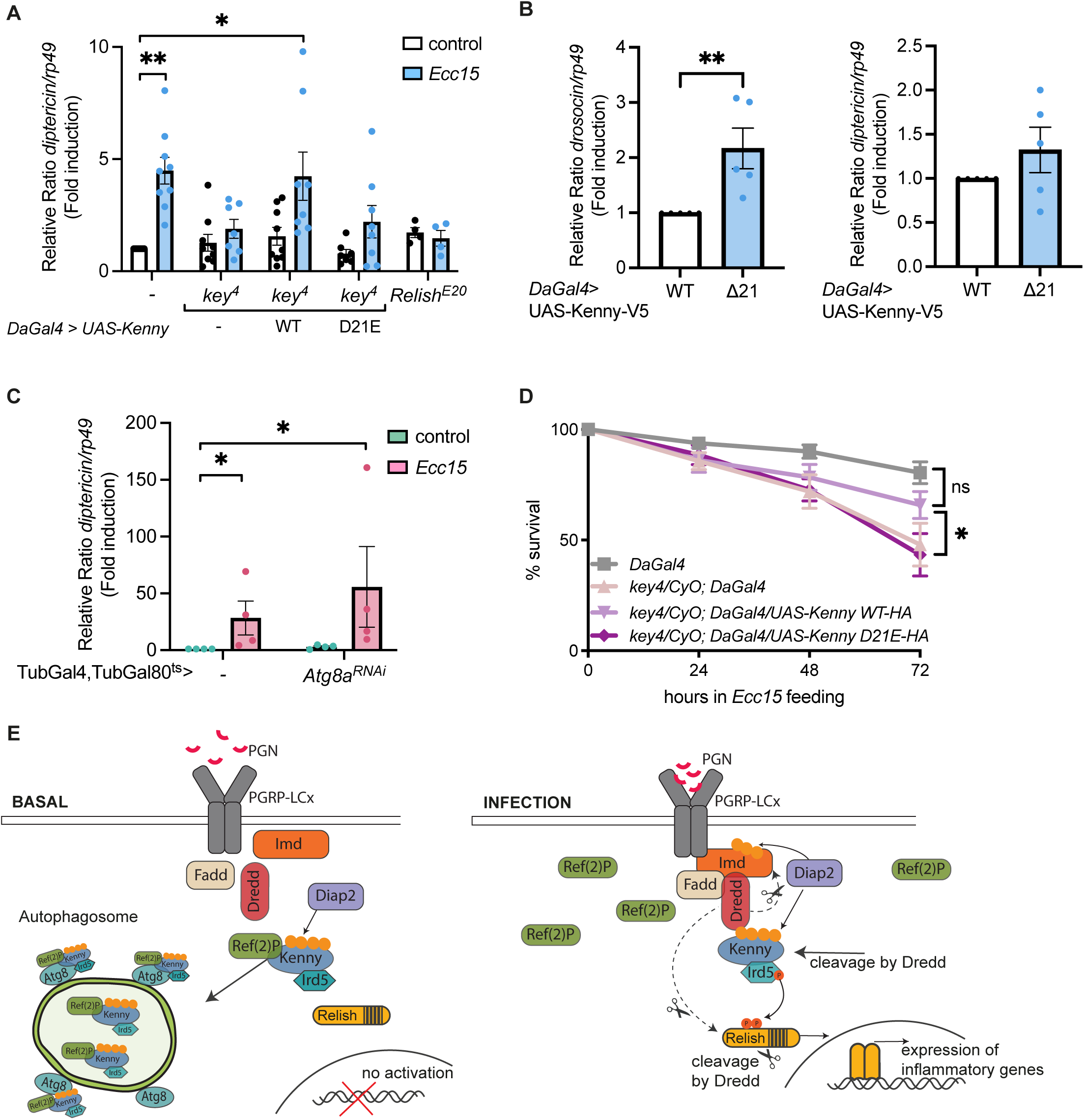
Kenny cleavage is crucial for NF-κB activation and intestinal immune responses. **(A)** HA-tagged wildtype or D21E mutant Kenny were expressed under the control of *DaGal4* in a *key^4^*/*CyO* Kenny mutant heterozygous background. Adult control *DaGal4* flies, *key^4^*/*CyO* files, the Kenny expressing flies, and Relish mutant flies were fed with 5% sucrose and *Ecc15* for 16 h and guts were dissected. Relish activation was studied by analysing the expression of *diptericin* with qPCR. Error bars indicate SEM from independent experimental repeats using at least 8 guts per repeat, n>3. **(B)** V5-tagged wildtype or Δ21 ubiquitin-fusion mutant Kenny were expressed under the control of *DaGal4.* Adult flies were fed with 5% sucrose and *Ecc15* for 16 h and guts were dissected. Relish activation was studied by analysing the expression of *drosocin* and *diptericin* with qPCR, n>3. Error bars indicate SEM from independent experimental repeats using at least 8 guts per repeat. **(C)** Adult control TubGal4, TubGal80^ts^ flies and flies expressing Atg8a^RNAi^ under control of TubGal4, TubGal80^ts^ were fed with 5% sucrose and *Ecc15* for 16 h. Relish activation was studied by analysing the expression of *diptericin* with qPCR. Error bars indicate SEM from independent experimental repeats using at least 8 adult flies per repeat, n=4. **(D)** HA-tagged wildtype or D21E mutant Kenny were expressed under the control of *DaGal4* in a *key^4^*/*CyO* Kenny mutant heterozygous background. Adult control *DaGal4* flies, *key^4^*/*CyO* files, and the Kenny expressing flies were fed with 5% sucrose and *Ecc15* and their survival was monitored over time, n>3. Error bars indicate SEM from independent experimental repeats using 20 flies per repeat. **(E)** Schematic representation of the proposed model: under basal conditions, Kenny localises to autophagosomes through its association with Ref(2)P and Atg8a. Upon infection, Dredd becomes activated and cleaves Imd, Relish, and Kenny. This cleavage event separates Kenny’s LIR domain, resulting in Kenny stabilisation and Ref(2)P accumulation, promoting efficient Imd signalling.

To study if the loss of NF-κB activation in Kenny mutant flies translated to a loss of immunity, we studied whether the cleavage of Kenny is crucial for flies to survive bacterial infection by feeding the pathogen *Ecc15*. Control flies can mount an immune response and survive the infection, whereas the Kenny LOF *key^4^*-mutant flies were sensitive to bacterial infection. As expected, we found that transgenic expression of wildtype Kenny rescues the Kenny LOF phenotype, as they survive *Ecc15*-infection. The expression of cleavage-resistant Kenny does however not rescue the Kenny LOF phenotype, suggesting that non-cleaved Kenny is not able to function in Imd signalling (Figure 5D).

## Discussion

In *Drosophila*, inflammatory signalling pathways such as the NF-κB-activating Imd pathway are activated in response to infection induced by pathogenic bacteria^6,10^. However, this activation needs to be especially tightly restrained in cells with a barrier function, such as the intestinal epithelium that exists in an environment of commensal bacteria. To prevent constitutive activation of the Imd pathway, the IKK complex is degraded via autophagy^26^. Here, we show that the selective autophagy receptor Ref(2)P interacts with Kenny to target the IKK complex to autophagosomes for degradation, thereby preventing Imd pathway activation. Upon Imd activation Dredd binds Kenny and disrupts its interaction with Ref(2)P leading to accumulation of Ref(2)P (Figure 5E). This signal-induced interaction depends on catalytical activation of Dredd, as Kenny cleavage leads to separation of the LIR motif of Kenny diverting the IKK complex from autophagic degradation to enable it to function as an activator of the Imd pathway during intestinal pathogen exposure.

Autophagy has been shown to have a role in dampening NF-κB pathways by removing key signal transduction components^26^. In addition to the IKK complex, several other NF-κB regulating proteins, including the TAB/TAK complex, have been shown to interact with the autophagosome, either via own LIR motives or by association to LIR containing proteins^35^. Hence, it is possible that Dredd regulates the autophagosome-association of other proteins as well. The mammalian Kenny homologue NEMO does not contain a LIR domain of its own, but the related protein optineurin is a LIR-containing autophagy receptor^26,36^. Optineurin is an important regulator of inflammatory NF-κB and interferon signalling, and it has been shown to interact with many conserved autophagy-associated proteins^37^. It would be interesting to study if caspase-mediated cleavage regulates optineurin stability and function. In addition, caspases may also be able to separate other signalling sequences regulating protein stability, localisation or activity, both during inflammation and other conditions.

In this study, we suggest a new role for caspases in regulating the stability of proteins involved in cell death and survival decisions. Caspase-dependent cleavage of the LIR motif provides a mechanism for stabilising the *Drosophila* IKK complex by preventing Ref(2)P-dependent autophagy. It is also possible that Dredd activation itself is enhanced upon interaction with Kenny. It was recently shown that Caspase-8 cleaves human sequestosome-1 (p62), leading to stabilisation of p62, enhanced activity of Caspase-8, and sensitivity to TNF-induced cell death. In mammals, p62 can also directly interact with NEMO and functions as an adaptor linking ubiquitination-dependent signalling to NF-κB activation^47,51^. Mechanistically similarly to our findings in *Drosophila*, caspase-8 ubiquitination diverts TRAIL signalling to NF-κB activation by recruiting NEMO to a caspase-8 containing complex via the NEMO UBAN^38,40^. Consistent with this, Optineurin, which has strong homology to NEMO^41^, has been shown to interact with the prodomain of caspase-8, and this has been suggested to restrain apoptosis induced via TNF-R signalling complex II^32^. Likewise, interaction between NEMO and viral FLIPs, which contain DEDs similar to caspase-8 and Dredd, enhances viability by promoting activation of NF-κB signaling^33,34^. Taken together, we report an evolutionary conserved model in which caspase-IKK interactions and selective autophagy receptors cooperatively balance inflammatory signalling and cell survival decisions.

## Acknowledgements

We thank Neal Silverman for antibodies, Pascal Meier, François Leulier, Bloomington *Drosophila* Stock Center (NIH P40OD018537) and Vienna *Drosophila* Resource Center (VDRC) for fly stocks used in this study. Ida Bäckström and Christa Kietz are acknowledged for technical assistance. This study was facilitated by the Åbo Akademi University Fly Unit supported by Biocenter Finland. Imaging was performed at the Advanced Imaging Core Facility at Turku Bioscience Centre, supported by Biocentre Finland, the Finnish Advanced Microscopy Node of Euro-BioImaging Finland (Turku, Finland), and Turku Bioimaging (Research Council of Finland, FIRI, #359073, #358879, #367582 and #367577) and the HSØ Advanced Light Microscopy Core Facility at Oslo University Hospital. The research was funded by Research Council of Finland Project funding (#321850), the InFLAMES Flagship Programme of the Research Council of Finland (#337531), the Research Council of Finland strategic research profiling area Solutions for Health at Åbo Akademi University (#336355), the Sigrid Jusélius Foundation, the Swedish Cultural Foundation, the Magnus Ehrnrooth Foundation, Turku Doctoral Network in Molecular Biosciences, and the Center of Excellence in Cellular Mechanostasis at Åbo Akademi University.

## Author contributions

AKM and AMD contributed to the design, execution and analysis of most of the experiments and writing of the manuscript. ALA performed transfections of S2 cells, Western blotting of fly lysates and qPCR. KK performed qPCR, HA and GFP-immunoprecipitations. VL performed dissections and Western blotting of fly guts and survival analysis. FS performed confocal imaging on whole fly guts and HA and GFP-immunoprecipitations. EH performed transfections and, Western blotting of S2 cell lysates and cloned constructs for generation of transgenic flies. TER contributed reagents, *Drosophila* strains and scientific expertise. AM contributed to experimental design, data analysis, structural modelling and writing of this manuscript.

## Materials and methods

### Fly husbandry

*Drosophila melanogaster* were maintained at 25°C on Nutri-fly BF (Dutscher Scientific, Essex, UK) or on medium containing agar 0.6% (w/v), malt 6.5% (w/v), semolina 3.2% (w/v), baker’s yeast 1.8% (w/v), nipagin 2.4%, and propionic acid 0.7% (Hi-Fly University of Helsinki Drosophila core facility, Helsinki Institute of Life Science) in a 12 h light–dark cycle. Adult flies were used for all experiments described. Wildtype *Canton^S^* or driver lines were used as controls. *Canton^S^,* the driver line *DaughterlessGal4 (DaGal4),* balancer lines, *UAS*LJ*Dredd, dredd^D44^* (with a G98R mutation), *dredd^L23^* (with a W436R mutation)^11^ and *key^4^/CyO*^17^ fly lines were kindly provided by Prof. Pascal Meier. The *NP1Gal4* driver line was provided by Prof. François Leulier. The *TubGal4; UAS-mCherry-Ref(2)P* from Assoc. Prof. Tor Erik Rusten and genomic *3xmCherry-Atg8a* was kindly provided by Professor Gabor Juhasz. The *w*; P{UASp-EGFP-key}3* (stock #90309, referred to as *UAS-GFP-Kenny)* and w*; P*{UAS-Atg8a-GFP}2* (stock #52005) were obtained from Bloomington stock centre. The *UAS-Atg8a^RNAi^*(stock #109654) was obtained from Vienna *Drosophila* Resource Center (VDRC). Fly egg injection for generation of HA- or V5-tagged UAS-Kenny transgenic flies was done by Bestgene Inc. HA-tagged UAS-Kenny^WT^ and UAS-Kenny^D21E^ and V5-tagged ubiquitin fusion wildtype and Δ21 mutant Kenny were introduced to the landing site line #24749^43^, and expression of the transgenes was verified by Western blotting for HA-tag and V5-tag.

### Plasmids and antibodies

Plasmids pMT/Flag-His, pMT/HA-Flag, pMT-Dredd-HA, pMT-Dredd-V5, and pMT-V5-Kenny-V5, were kindly provided by Prof. Pascal Meier. Kenny-HA was subcloned from pMT-V5-Kenny-V5. Site-directed mutagenesis to make Kenny D21E, D27E, D67E, D88E mutants was performed using QuikChange Lightning Site-directed Mutagenesis Kit (Agilent Technologies). HA-tagged UAS-Kenny^WT^ and UAS-Kenny^D21E^ were subcloned from pMT-Kenny-HA to a pUASTattB plasmid for injection to fly eggs. A constitutively cleaved Δ21 mutant Kenny was made using a ubiquitin fusion strategy that enables expression of neo-N-termini proteins by translational processing^48^. A corresponding ubiquitin fused wildtype Kenny construct was generated as an expression control. The following antibodies were used: anti-GFP (ab6556, Abcam), anti-HA (clone 3F10, #11867423001, Roche), anti-V5 (Clone SV5-Pk1, #MCA1360, Bio-Rad), anti-Ref(2)P (ab178440, Abcam), anti-Ref(2)P, and anti-Actin (C-11, sc-1615, Santa Cruz). Anti-Kenny (IKKγ 1926) was kindly provided by Neil Silverman.

### Cell culture

*Drosophila* Schneider S2 cells (Invitrogen) were grown at 25□°C using Schneider medium supplemented with 10% fetal bovine serum, 1% l-glutamine, and 0.5% penicillin/streptomycin. Effectene transfection reagent (Qiagen) was used to transfect indicated constructs according to manufacturer’s instructions. Expression of construct in pMT plasmids was induced using 0.5 mM CuSO_4_ for 16 h before lysis.

### Bacterial strains and infection experiments

The Gram-negative bacteria *Erwinia carotovora carotovora 15* (*Ecc15*), kindly provided by Dr. François Leulier, was cultivated in Luria-Bertani (LB) medium at 29□°C for 16□h with continuous shaking and concentrated to a desired optical density (OD). For all Western blotting and qPCR analyses, as well as the survival assays OD 0.2 was used. For the survival in Figure 5C, OD 0.1, was used. For survival assays, 20 flies were counted at indicated time points after infection. Survival experiments, in which wildtype flies survived to a less extent than 30%, were excluded. These criteria were pre-established. For all Western blot and qPCR analyses infection experiments were performed by feeding adult flies for 16 h with a 1:1 solution of *Ecc15* and 5% sucrose added on a Whatman paper before dissecting guts. Treatment with inhibitors were performed by supplementing 100□µM chloroquine, diluted in a 1:1 solution of 5% sucrose and either H_2_O or *Ecc15*. The flies were then fed for 16 h before analysing their dissected guts by Western blotting and confocal imaging.

### Lysis of fly guts for Western blotting

Ten intestines from adult *Drosophila* were dissected, homogenized and lysed for 10□min on ice in a buffer containing 50□mM Tris (pH 7.5), 150□mM NaCl, 1% Triton X-100, 1□mM EDTA, 10% glycerol and Pierce™ Phosphatase inhibitor. The lysates were cleared before the addition of Laemmli sample buffer, and protein levels were analysed by Western blotting.

### HA, V5 and GFP immunoprecipitations

S2 cells and flies were lysed in a buffer containing 50□mM Tris (pH 7.5), 150□mM NaCl, 1% Triton X-100, 10% glycerol, 1□mM EDTA, 5□mM NEM, 5□mM chloroacetamide and Pierce™ Protease and phosphatase inhibitor. Cell lysates were cleared by centrifuging at 12,000□rpm for 10□min at 4□°C. The samples were incubated with anti-GFP agarose beads (ChromoTek) for 4 h or 16 h or anti-HA or anti-V5 agarose beads (Sigma) for 2□h under rotation at 4□°C. The beads were washed three times in a wash buffer containing 10□mM Tris (pH 7.5), 150□mM NaCl, 0.1% Triton X-100 and 5% glycerol. HA, V5 or GFP-conjugated proteins were eluted using Laemmli sample buffer.

### Quantitative RT-PCR (qPCR)

Eight intestines from adult *Drosophila* were homogenised using QIAshredder (Qiagen) and total RNA was extracted with RNeasy Mini Kit (Qiagen) and cDNA was synthesised with SensiFast cDNA synthesis kit (Bioline, London, UK) according to the manufacturers’ protocols. qPCR was performed using SensiFast SYBR Hi-ROX qPCR kit (Bioline). *rp49* was used as a housekeeping gene for ΔΔCt calculations. The following gene-specific primers were used to amplify cDNA: *diptericin* (5’-ACCGCAGTACCCACTCAATC, 5’- ACTTTCCAGCTCGGTTCTGA), *drosocin (5’-*CGTTTTCCTGCTGCTTGC, 5’-GGCAGCTTGAGTCAGGTGAT), and *rp49* (5’-GACGCTTCAAGGGACAGTATCTG, 5’-AAACGCGGTTCTGCATGAG).

### Immunofluorescence and imaging

*Drosophila* intestines were carefully dissected in PBS and immediately fixed in PBS containing 4% formaldehyde for 1 h at room temperature on a shaker. After fixation, the samples were washed two times with 1x PBS before incubation in blocking buffer (PBS containing 0.3% Triton X-100 and 0.3% BSA) for 1 h. Primary antibodies were incubated in a concentration of 1:200 or 1:600 in blocking buffer at 4□°C, anti-Ref(2)P overnight and anti-HA for 48 h. The intestines were washed three times with blocking solution, and secondary antibody stainings were done in blocking buffer overnight at 4□°C. Secondary antibodies were purchased from Jackson ImmunoResearch. The intestines were mounted in Vectashield with DAPI for imaging. Samples were imaged on Zeiss LSM880 confocal microscopes and Leica Stellaris 8 Falcon confocal microscope and further processed with FIJI^49^.

### Statistics

Results from survival assays were analysed by two-way analysis of variance (ANOVA) with Dunnett’s post hoc test for 95% confidence intervals. Results from qPCR were analysed by two-tailed unpaired t-test on relative fold induction of the target gene compared to a normalised control sample (-ΔΔCt). Normalised results from qPCR were analysed with Mann-Whitney U test when two samples were compared, Kruskal-Wallis test with Dunnett’s post hoc was applied when several samples were analysed. Relative protein expression from western blots was quantified with ImageJ and analysed by two-tailed Student’s t-test. In comparison to normalised control values, the Mann-Whitney U test was applied. Statistical analyses were performed using GraphPad Prism version 10.3.0 for Windows (GraphPad Software, San Diego, California, USA). In figures, ns stands for nonsignificant, * p < 0.05, ** p < 0.01, *** p < 0.001, **** p < 0.0001. Error bars in figures specify SEM from the indicated number of independent experimental repeats. Experiments were performed indicated number of times (n ≥ 3).□

