## Supplementary figures 1-4 for "Dredd-mediated cleavage of Kenny uncouples the IKK complex from selective autophagy to enable innate immunity"

Supplementary Figure 1

**Dredd-mediated cleavage of Kenny uncouples the IKK complex from selective autophagy to enable innate immunity**

Mohan et al.

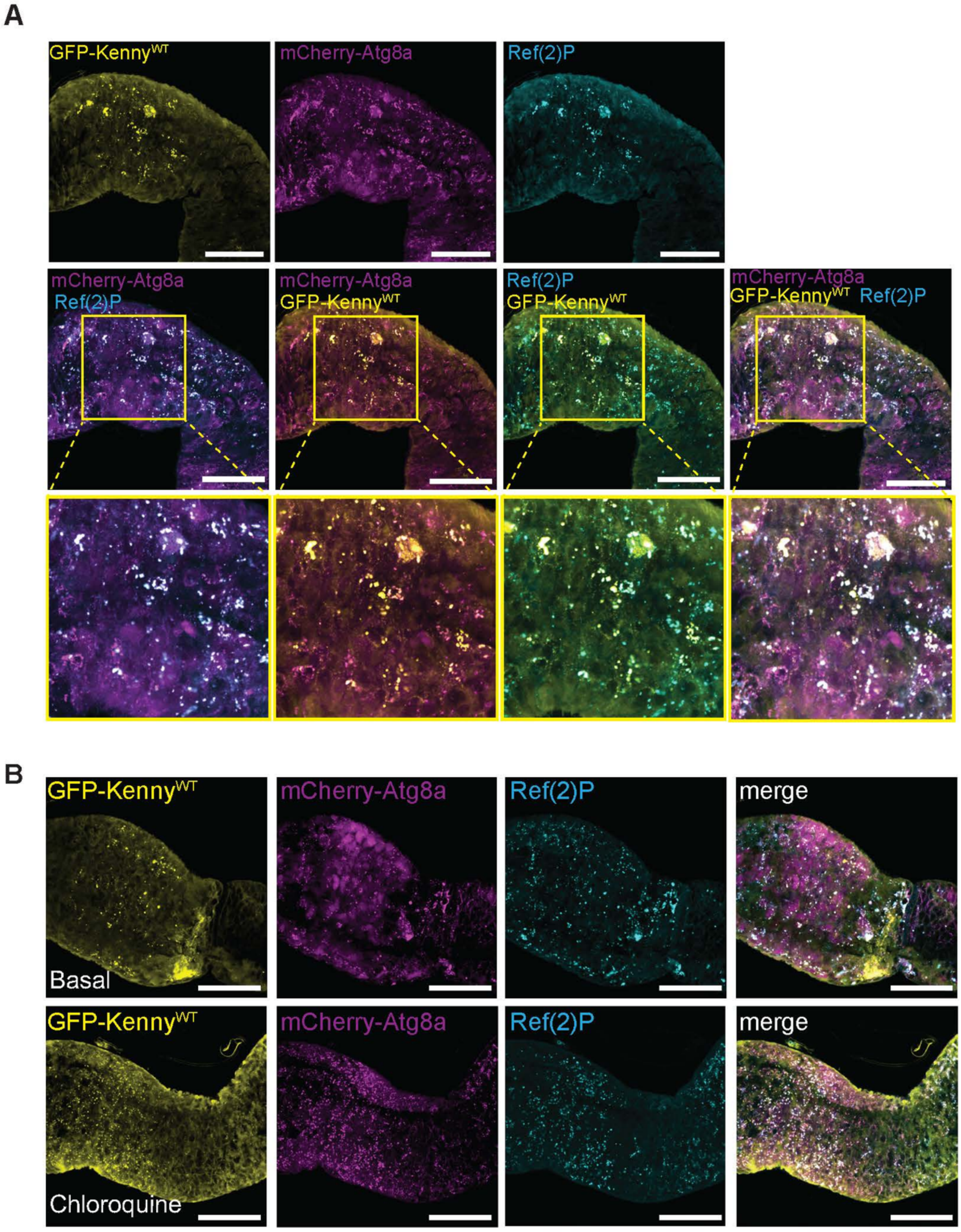


**Supplementary Figure 1. Kenny co-localisation with Ref(2)P and Atg8a is enhanced upon stabilisation by autophagosome inhibition. (A)** Dissected intestines from adult flies expressing GFP-Kenny (yellow) and mCherry-Atg8a (magenta) were stained for Ref(2)P (cyan). Midguts were imaged by confocal microscopy using at least 3 intestines per repeat, n=3, scale bar 100 µm. The upper panels show localization of Kenny, Atg8a and Ref(2)P separately. The middle panels show merge of Atg8a and Ref(2)P, Kenny and Atg8, Ref(2)P and Kenny, and all three channels together. The lover panels are zoom-in images of the squares in the middle panel. Co-localisation is visible as white puncta. **(B)** Adult flies expressing GFP-Kenny (green) and mCherry-Atg8a (red) were fed with 5% sucrose and 100 µM chloroquine for 16 h. Dissected intestines were stained for Ref(2)P (cyan). Midguts were imaged by confocal microscopy using at least 3 intestines per repeat, n=3, scale bar 100 µm. The images at the right are merges of all three channels.

Supplementary Figure 2 and 3

**Dredd-mediated cleavage of Kenny uncouples the IKK complex from selective autophagy to enable innate immunity**
Mohan et al.

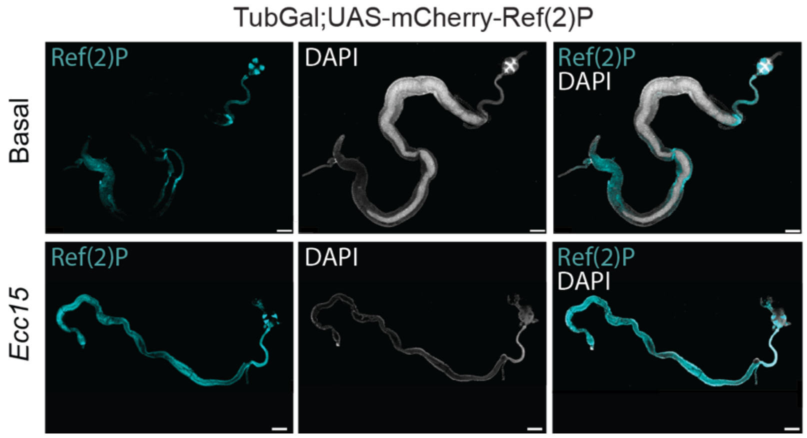


**Supplementary Figure 2. Bacterial infection stabilises Ref(2)P.** Adult TubGal;UAS-mCherry-Ref(2)P flies were fed with 5 % sucrose and *Ecc15* for 16 h. Dissected intestines were stained with DAPI to visualised nuclei and imaged by confocal microscopy. mCherry-Ref(2)P is indicated in cyan and DAPI in white. Scale bar 100 µm, n=3.

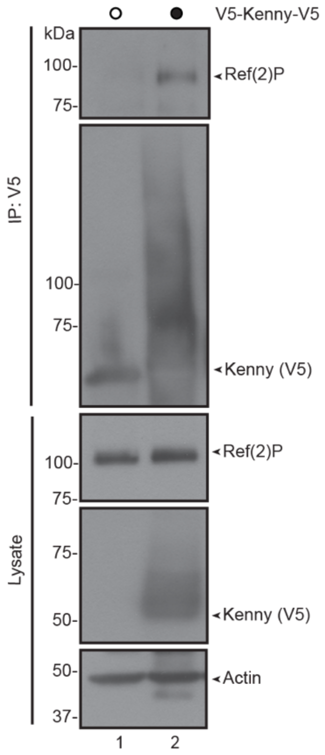


**Supplementary Figure 3. Ref(2)P and Kenny interact in S2 cells.** *Drosophila* S2 cells were transfected with empty vector or V5-tagged Kenny. V5-immunoprecipitations were performed and the samples were analysed by Western blotting with anti-Ref(2)P, anti-V5, and anti-Actin antibodies, n>3.

Supplementary Figure 4

**Dredd-mediated cleavage of Kenny uncouples the IKK complex from selective autophagy to enable innate immunity**

Mohan et al.

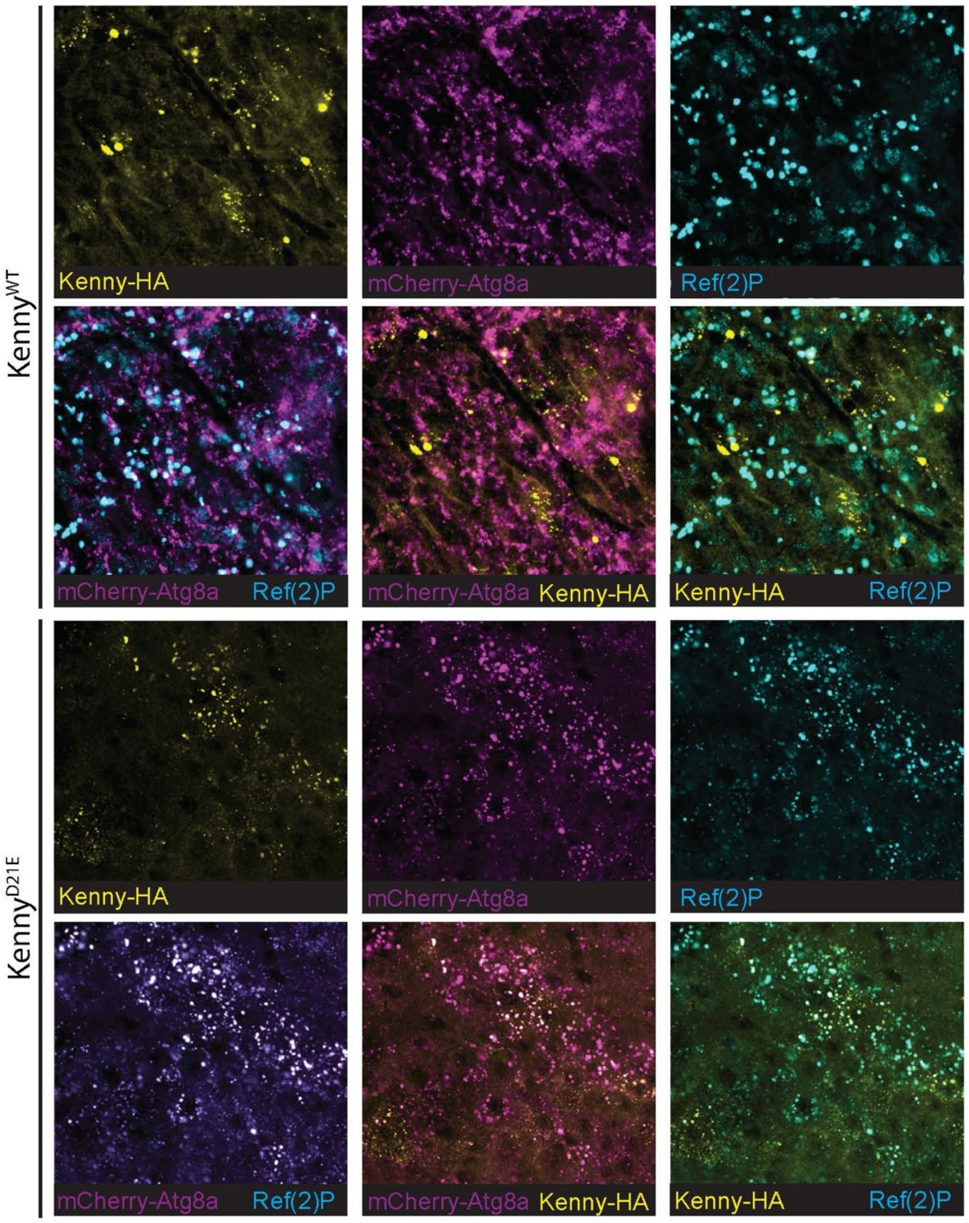


**Supplementary Figure 4. Infection stabilises Kenny against autophagic degradation.** Dissected intestines from adult flies expressing mCherry-Atg8a (red) crossed with flies expressing HA-tagged wildtype or D21E mutant Kenny were stained for HA (green) and Ref(2)P (magenta). Midguts were imaged by confocal microscopy using at least 3 intestines per repeat, n=3. Scale bar 100 µm. The first and third rows of panels show localization of Kenny, Atg8a and Ref(2)P separately. The second and fourth rows of panels show merge of Atg8a and Ref(2)P, Kenny and Atg8, Ref(2)P and Kenny, and all three channels together.
